# Chemical stressors differentially induce prophage replication in *Acinetobacter baumannii* strains

**DOI:** 10.64898/2026.09.15.751888

**Authors:** Jessica Trinh, Hans K. Carlson, Vivek K. Mutalik, Catherine M. Mageeney

## Abstract

Prophages are prevalent genomic elements that profoundly shape microbial communities by controlling host bacterial populations through lysis or enabling genetic recombination via transduction. Standard induction protocols rely heavily on DNA-damaging agents like mitomycin C, failing to capture the complex chemical landscapes bacteria encounter in natural and clinical settings where diverse environmental insults trigger excision. Furthermore, the regulatory variations governing polylysogenic hosts containing multiple co-habiting prophages under identical stressors remain largely uncharacterized. In this study, we surveyed a panel of five clinical *Acinetobacter baumannii* strains containing zero to four resident prophages against a library of 57 diverse chemical stressors. We demonstrate that chemical treatments, including distinct classes of antibiotics, drive differential prophage induction patterns across different host backgrounds. This could lead to a greater understanding of the importance of prophages in antibiotic treatment regimes.

**Importance:** Our study highlights the need for a greater understanding of phage replication during various treatment regimens and changes in the environment. Our simple, high-throughput assay allows us to examine strains containing multiple prophages, which have been underexplored. Additionally, the mechanisms allowing prophage activation in one strain versus another under the same stressor remain to be understood.

## Introduction

Bacteriophages are abundant and diverse within human microbiomes and have the potential to play a major role in human health (Shkoporov and Hill 2019, Liang and Bushman 2021). Phages in human microbiomes are predicted to infect several human-associated bacterial genera which can lead to lysis of their bacterial host or the transfer of genetic material through horizontal gene transfer mechanisms (Camarillo-Guerrero *et al*. 2021, Borodovich *et al*. 2022). However, the exact role that phages play in regulating and modulating the human microbiome is still not fully understood, especially for pathogens. There are two types of phages: lytic phages that actively replicate and lyse hosts, and temperate phages that integrate into host genomes as prophages (Lwoff 1953). An estimated 53.9% of all bacterial genomes contain at least one prophage (Yu *et al*. 2025).

While there is a sizeable amount of literature examining the induction of a single prophage within a bacterial strain, there are still many unknowns around polylysogens (bacteria with multiple prophages). It is possible that in a single host, prophages experience within-host competition for resources, leading to the evolution of alternative mechanisms for induction, responding to different induction cues, or impacting the excision and/or replication of other prophages by modulating bacterial gene expression (Silpe *et al*. 2023, Azulay *et al*. 2022).

The most well-studied prophage inducer is DNA damage, particularly by mitomycin C (MMC), (DeMarini and Lawrence 1992, Silpe *et al*. 2023a). DNA damage induces prophage excision by activating the SOS response, a conserved bacterial signaling pathway (Maslowska *et al*. 2018, Silpe *et al*. 2023b). This response triggers the self-cleavage of the phage repressor protein in many different prophages (Thabet *et al*. 2023, Atsumi and Little 2006, Geslewitz *et al*. 2025). However, there are other inducers of excision such as heat, antibiotics, and bile salts, which correlate with other bacterial stress pathways (Armentrout and Rutberg 1971, Henrot and Petit 2022). Other classes of compounds cause bacterial stress and likely induce prophages as well (Hu *et al*. 2026, Carlson *et al*. 2018, Guan *et al*. 2017). Treatment of bacteria with metals can cause redox stress (Cuypers *et al*. 2010, Alquethamy *et al*. 2021), depletion of essential metal ions from enzymes (Osorio-Rico *et al*. 2017, Pal *et al*. 2022), and disruption of lipid membranes (Warnes *et al*. 2011, Quaranta *et al*. 2011). Compounds that can generate reactive oxygen species (ROS), such as hydrogen peroxide, can also cause microbial stress through DNA damage and the oxidation of metabolic enzymes (McDonnell and Russell 1999, Imlay 2013). In addition to targeting specific cellular processes, antibiotics trigger off-target effects through ROS production and DNA damage (Kohanski *et al*. 2010), which also contribute to bacterial stress.

Phage-antibiotic synergy (PAS) has been leveraged as a potential treatment strategy for multidrug-resistant bacteria in recent years (Zhao *et al*. 2024, Liu *et al*. 2020, Ryan *et al*. 2012). By treating a patient with sub-lethal concentrations of antibiotics alongside the phage, these treatments have shown an increase in phage penetration of biofilms (Kumaran *et al*. 2018, Segall *et al*. 2019), and the re-sensitization of some phage-resistant strains to antibiotics (Ho *et al*. 2018, Chan *et al*. 2016). While antibiotics are known inducers of prophage excision (Henrot and Petit 2022), whether the resulting phage particles also contribute to reducing pathogen populations depends on the genomic content of the prophage (Gavric and Knezevic 2026, Cook and Hynes 2025). PAS has resulted in unexpected outcomes such as when prophage induction triggers the upregulation of toxin genes in *Clostridium dificile* (Govind *et al*. 2009, Sekulovic *et al*. 2011) and promoting *Enterococcus faecalis* virulence through increased adhesion to platelets (Matos *et al*. 2013). Recent studies show synergy between sub-lethal antibiotic treatments and prophages, suggesting that prophages play a direct role in successful treatment outcomes (Fatima *et al*. 2025, Al-Anany *et al*. 2021).

In this paper, we evaluated a wide range of bacterial stress compounds for a thorough survey of prophage inducers in *Acinetobacter baumannii*. This species is a clinical bacterial pathogen on the World Health Organization list of drug-resistant bacteria that pose the greatest threat to human health (Willyard 2017). *A. baumannii* strains have been predicted to contain multiple prophages, which often contain virulence factors that may be important to *A. baumannii* colonization and persistence in human hosts (Costa *et al*. 2018, Trinh *et al. 2026*). Understanding specific chemical triggers of prophage excision in a relevant human pathogen could provide insights into how prophages behave in different environments and under antimicrobial treatments. This knowledge could help researchers leverage prophage induction for improved treatment options for bacterial infections.

## Results and Discussion

### MRSN strains have differential growth in response to different microbial stressors

We leveraged a previously developed library of stressors, which includes metals, antibiotics, and salts (Price *et al*. 2018, Hu *et al*. 2026, Carlson *et al*. 2017, https://fit.genomics.lbl.gov/), to screen a set of *A. baumannii* strains from the Multidrug-Resistant Organism Repository and Surveillance Network (MRSN) for prophage induction. Concentrations required to reduce bacterial populations by half (IC50s) for stressors were determined across four MRSN strains in 384-well plates (**Supplemental Table S1**). For all subsequent experiments we used the IC50 concentration determined from experiments with MRSN31468 to stress strains by diluting from concentrated stocks (**Supplemental Table S2**). Of 79 initial stressors, 57 were selected for further experiments based on assay constraints and whether the stressor caused high enough levels of growth inhibition to feasibly facilitate prophage replication.

Our overall goal is to understand how prophages respond to a wide variety of stressors. We used our TIGER/Islander pipeline (Mageeney *et al*. 2020b) to predict prophage in four *A. baumannii* MRSN strains (**Supplemental Table S3**). We selected a set of strains that contained 0 – 4 prophages that ideally contain a full phage gene complement, are in the 30 to 60 kbp range, and are in unique prophage clusters (**Table 1**). We used the precise attachment sites (*att*) generated by the software to design primers for a qPCR-based assay we developed to quantify bacteriophage only when actively excising and replicating (Waggoner *et al*. 1974, Mageeney *et al*. 2020b).

**Table 1.**
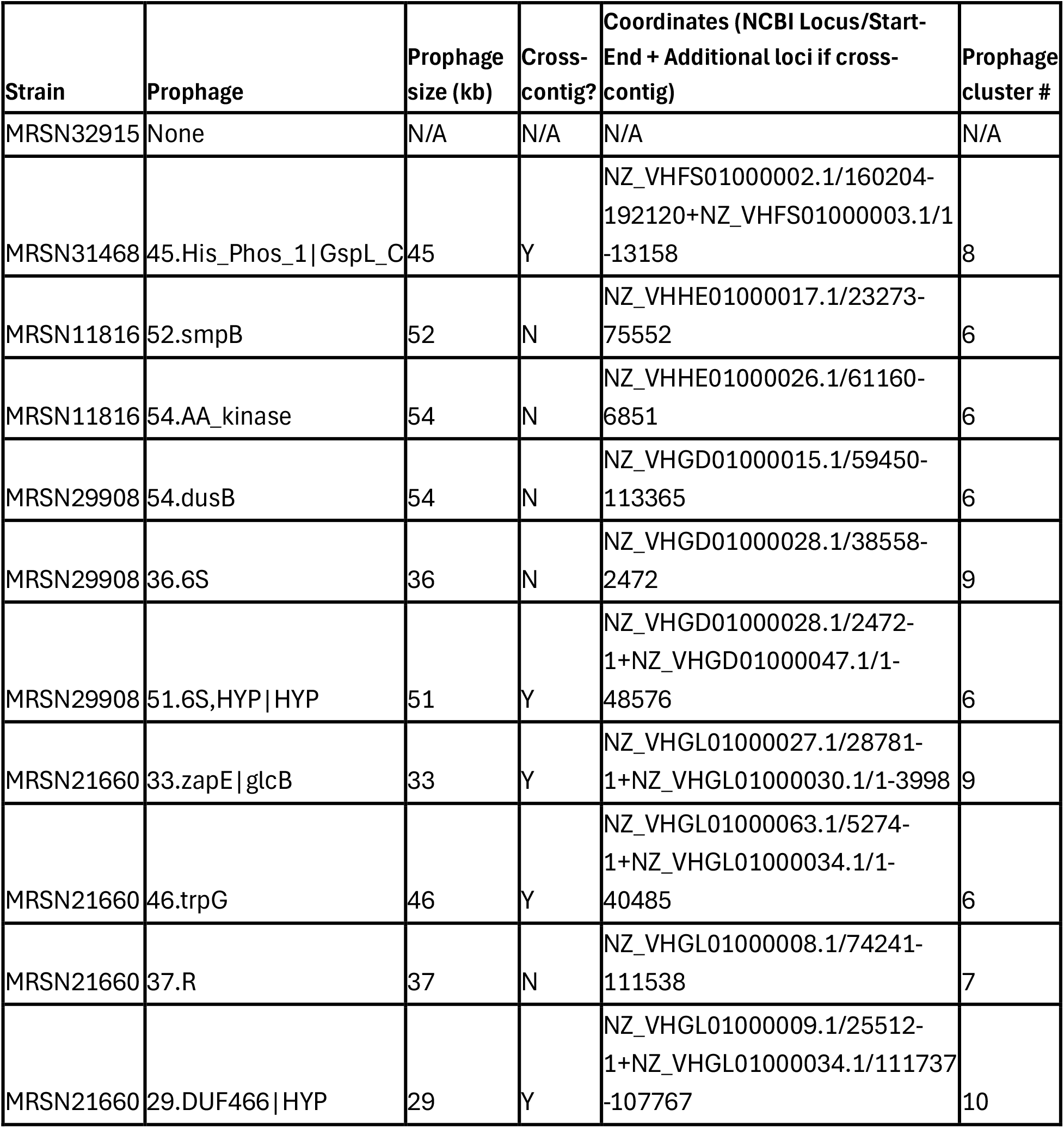
Strains selected for prophage screening, along with their corresponding prophages and prophage coordinates. Prophages are named by the size of the prophage in kilobases and their integration site. Clusters are indicated in **Supplemental Figure S3**.

Each strain was grown for 24 hours in LB in 96-well plates amended with the MRSN31468 IC50 concentration of stressor determined in the prescreens. Overall, we found that MRSN29908 and MRSN21660, which contain three and four prophages respectively, tend to be more resistant to a wider variety of stressor compounds compared to the other strains (**Figure 1**). At the concentrations tested, we did observe resistance of MRSN29908 and MRSN21660 to multiple antibiotics, which is consistent with literature observations (Galac *et al*. 2020, **Supplemental Table S4**). Interestingly, the strain containing no predicted prophage (MRSN32915) was similarly sensitive to many of the stressors as MRSN31468 and MRSN11816, which contain one and two prophages respectively. Conversely, MRSN29908 and MRSN21660 were more tolerant of many of the antibiotics, ions, and metal stressors used, particularly for the manganese (II) chloride, sodium chlorite, and magnesium chloride treatments. All of the tested strains grew to similar ODs as untreated controls at 24-hours when treated with metals, including zinc, nickel, or copper, which are known to display antimicrobial properties at high concentrations (Hassan *et al*. 2017, Khan *et al*. 2013, Williams *et al*. 2016). Some stressors, such as thallium (I) acetate, cadmium chloride, and apramycin, inhibited the growth of all the strains tested under these conditions (**Figure 1**). It is important to note that these results capture inhibition at a single 24-hour timepoint. Some strains may have been inhibited earlier but recovered by 24 hours, and strains inhibited after 24 hours may recover at later time points.

**Figure 1.**
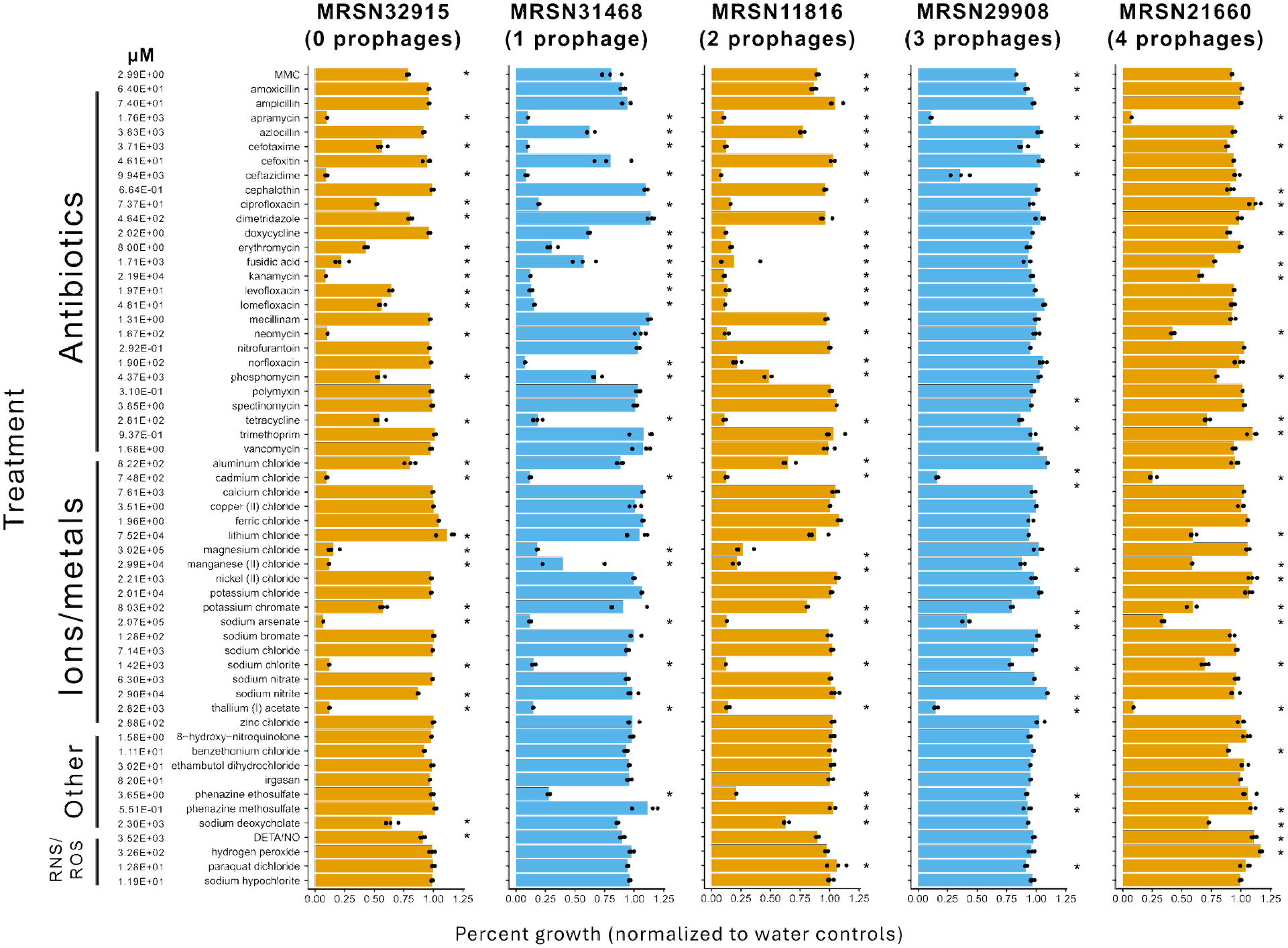
Growth data for each of the selected strains in various stressors. Each data point represents one well, and all treatments are performed in triplicate. Data are presented as percent growth, where percent growth = final OD_treatment_ / final OD_average_water_ * 100%. Asterisks (*) represent significantly different (p < 0.01) percent growth compared to water controls. The concentration of each stressor is listed in µM.

### Prophage excision is observed with a diverse set of compounds

We harvested, filter-sterilized, and normalized all lysates according to final ODs. We then DNase-treated supernatants to quantify relative amounts of excised prophage using qPCR. Treatments that greatly inhibited bacterial growth were excluded from qPCR analysis, it is likely that cells which are unable to grow in a given treatment are also unable to support prophage replication. We used water treatments as a baseline for spontaneous induction and MMC as a positive control. In general, treatments that inhibit strain growth by more than 50% tend to produce lower –dCt values with some exceptions (**Supplemental Figure S1**), suggesting a correlation between bacterial growth and prophage replication.

Antibiotics can be weak inducers in certain strains and induce prophage in others (**Figure 2**). Specifically, in MRSN31468 and MRSN11816, amoxicillin, ampicillin, and azlocillin induce prophage excision but they did not in MRSN29908 and MRSN21660, despite all strains showing similar growth relative to controls at 24 hours. Ceftazidime was only able to induce prophage replication in MRSN29908 and completely inhibited the growth of susceptible strains MRSN31468 and MRSN11816. In MRSN29908 ceftazidime also significantly inhibited growth despite the strain being documented as resistant, which may suggest that under these conditions, induced prophages are lysing MRSN29908 cells.

**Figure 2.**
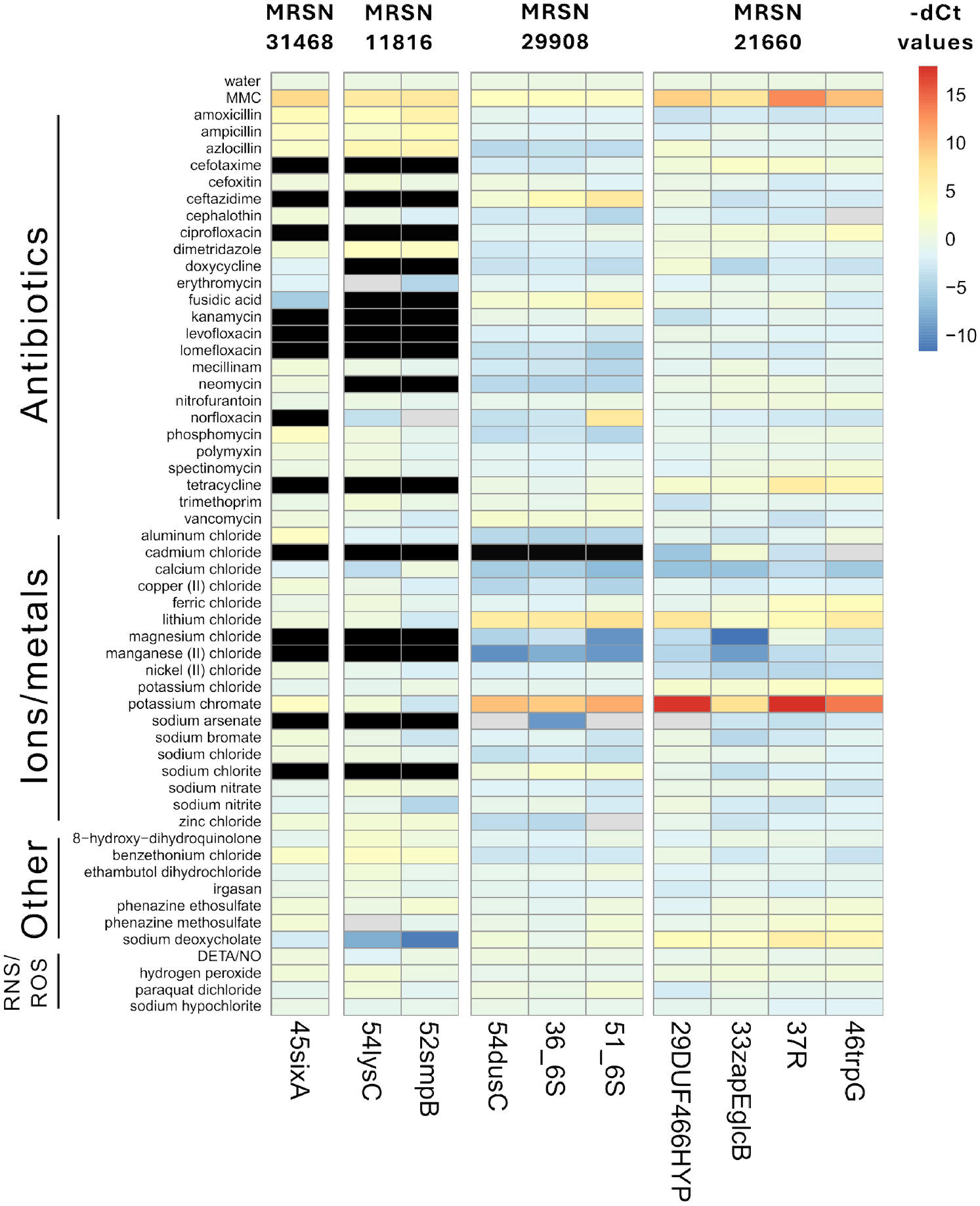
Heatmap of -ΔCt values for each prophage from their respective host, where - ΔCt = –(ΔCt_treatment_average_ – ΔCt_water_average_). Treatments are categorized broadly and more specific compound types can be found in **Supplemental Table S1**. Values above 0 indicate prophage that are induced compared to water. Black blocks indicate treatments that were excluded from further screening because the strain’s growth was severely inhibited (<20% growth). Gray blocks indicate treatments where no Ct values could be determined from the sample.

MRSN11816 and MRSN31468 were sensitive to four antibiotics tested in our screen, including ciprofloxacin, kanamycin, levofloxacin, and lomefloxacin, and were unable to grow in those antibiotics, meaning, the link between these antibiotics and prophage induction could not be evaluated. However, the other two prophage-laden strains (MRSN29908 and MRSN21660) were resistant to or tolerant of these antibiotics but showed a neutral or lower level of prophage induction compared to water. It is possible that a lower dose of the antibiotics for the more sensitive strains, we would have induced enough cell stress for prophage replication but not for complete inhibition of growth. It is also possible that the prophages contributed to the complete inhibition of these strains in some antibiotic treatments, which is another reason to test a lower dose of antibiotic. In the resistant or tolerant strains, these antibiotic compounds could also be unable to get into the cell or are removed from the cell rapidly, preventing cell stress. This observation highlights the need to carefully choose antibiotics to use to trigger the prophages in these strains and contribute to pathogen clearance. Prophage induction by antibiotics has been documented previously (Nair *et al*. 2025, Henrot and Petit 2022). While not all antibiotics directly affect DNA replication to cause DNA damage, antibiotics can also trigger the SOS response by affecting cell wall synthesis and triggering ROS production (Maiques *et al*. 2006, Miller *et al*. 2004, Gavric and Knezevic 2026, Baharoglu *et al*. 2013). Future experiments should determine the mechanisms behind prophage induction to better understand the synergy between antibiotic susceptibility and prophage replication. Further, it would be interesting to know if differences in prophage abundance between strains are a result of differential expression of SOS response genes or through another mechanism.

Out of the metal stressors we tested, we observed the strongest induction of prophage excision by potassium chromate in MRSN29908 and MRSN21660, though the magnitude of excision varied between individual prophages. Even though potassium chromate causes DNA damage and oxidative stress (Giacomucci *et al*. 2022, Pradhan *et al*. 2016), no prophage from MRSN31468 nor MRSN11816 seem to excise with the same treatment. For calcium chloride and manganese (II) chloride, we detected lower levels of prophage excision than in our water control, indicating an inhibitory effect of the treatment on prophage replication independent of bacterial growth. Phages often require divalent cation supplementation from Mg^2+^ or Ca^2+^ for efficient infection (Beumer *et al*. 1957, Bandara *et al*. 2012), however our results suggest that at a high enough concentration, these cations might prevent prophage excision or replication. We have also considered the effect of excess cations on PCR efficiency, since calcium ions are known to inhibit polymerase activity (Opel *et al*. 2010). We PCR-tested non-normalized lysate and found that the same product amplifies similarly in our magnesium and calcium treatments (**Supplemental Figure S2**). Surprisingly, none of the reactive oxygen species (ROS) stressors induce prophages, which was unexpected as ROS usually triggers the SOS pathway and has previously been shown to induce prophages (Braetz et al. 2026, Jancheva et al. 2025). It is possible that a higher concentration of ROS is required to induce prophage replication, considering most of the ROS treatments in our dataset do not significantly impede bacterial growth (**Figure 1, Supplemental Table S6**).

We wanted to determine whether similar prophages exhibit similar induction profiles. To do this, we first clustered prophages in our dataset based on their average amino acid identity and minimum percent of genes shared (Nayfach *et al*. 2021).

Prophages across three different strains are in the same cluster, which includes 54.lysC, 52.smpB, 54.dusC, 51.6S, and 46trpG (**Table 1, Supplemental Figure S3**). However, whether a prophage is able to excise and replicate in response to a particular treatment seems to be associated more within a strain rather than within a cluster. For example, sodium deoxycholate seems to inhibit prophage excision in MRSN11816’s 54lysC and 52smpB but is only either neutral or slightly inducing in the other prophages. This indicates that the bacterial stress response could play a greater role in prophage induction than prophage genetics alone. Prophages within the same strain have differing magnitudes of induction, as demonstrated by the differences in -ΔCt values for the four prophages in MRSN21660 when exposed to MMC. In MRSN29908, norfloxacin induces prophage 51.6S but not 36.6S or 54.dusC. Lithium chloride is a strong inducer of 29DUF466HYP and 46trpG excision but only a mild inducer for 37R in MRSN21660 (**Figure 2**). While it is difficult to assess whether prophages are consistently outcompeting other prophages across the entire population of cells, competition among prophages within the same host is a well-documented phenomenon (Silpe *et al*. 2013a, Sargen and Helaine 2024, Refardt 2011) and we have now demonstrated differential induction of prophages in the same host by chemical stressors.

## Conclusions

Here we used a high throughput screen to understand how various chemical stressor compounds impact prophage induction in *A. baumannii*. We observed prophage induction by diverse classes of stressors including antibiotics, inorganic ions and reactive oxygen and nitrogen species, with potassium chromate as the strongest inducer of prophage in two different *A. baumannii* strains. Strain background appeared to have a larger effect on prophage induction than stressor type or prophage relatedness in our experiments. Further research on prophage competition in response to inducers within a host or phage-antibiotic synergy could allow researchers to specifically target the induction of highly active prophages to aid in treatment options. Overall, we were able to discover new prophage inducers and possible prophage inhibitors that will can help to inform treatment of resistant bacterial infections.

## Materials and Methods

### Prophage prediction and clustering

Prophage prediction for *Acinetobacter baumannii* was performed as described previously (Mageeney *et al*. 2020b). Briefly, we used two platforms to predict prophages: Islander (Hudson *et al*. 2015), which looks for genomic islands in tRNA and tmRNA genes, and TIGER (Mageeney *et al*. 2020a), which looks for integrase-containing genomic islands. This analysis was performed on a diverse reference panel of 100 *A. baumannii* strains from the Multidrug-Resistant Organism Repository and Surveillance Network (MRSN) collection (Galac *et al*. 2020). The resulting genomic islands were annotated by our pipeline’s Tater software and sorted into categories with a custom script based on whether islands contained a credibly complete set of phage genes (Mageeney *et al*. 2020a). Plausible prophage sequences were clustered by the Markov Cluster Algorithm based on an average amino acid identity (>40%) and minimum percent of genes shared (>20%, Nayfach *et al*. 2021). One strain was selected for each different quantity of prophage, from zero to four prophages, capturing as many phages in different clusters as possible.

### Initial screening of Acinetobacter baumannii strains with stressors

For high-throughput dose–response assays, concentrated stock solutions were serially-diluted across three 384-well plates (Costar) using a Biomek FxP liquid handling robot (Beckman Coulter) to attain 12 two-fold dilutions of each compound (Carlson *et al*. 2023, Day *et al*. 2024, Hu *et al*. 2026). These arrayed dose–response plates were inoculated 1:1 with 2X concentrated LB medium with a suspension of each *A. baumannii* strain at an initial OD600 of 0.02. Plates were sealed with breathable seals (BreathEasy, Sigma-Aldrich) and incubated aerobically at 30 °C. At defined time points, OD600 was measured on a Tecan M1000 Pro microplate reader (Tecan), and plate reads were uploaded to an in-house growth database. Early stationary-phase time points were selected for dose– response analysis, and custom scripts were used to fit a nonlinear regression to determine half-maximal inhibitory concentrations (IC50) for each compound against each strain (Carlson *et al*. 2023, 2019). IC50s for MRSN31468 were chosen for further screening with the other strains.

### Prophage induction and lysate harvesting in 96-well plates

Strains were inoculated in LB broth at 37 °C and grown overnight before induction assays. Strains were back-diluted 1:100 in 4 mL LB broth and grown to an OD of 0.4 – 0.6 before using the culture for microplate assays. Strains were back-diluted in 150 µL in 96-well microplates (Fisher Scientific #08-772-54) by adding 50 µL culture, 50 µL 2X concentrated LB broth, and 50 µL of stressor. All stressors were back-diluted to the MIC of MRSN31468 if possible, with some exceptions (**Supplemental Table S2**). Three wells were used for each treatment, and each plate included both a positive control (1 µg/mL mitomycin C, MMC) and a negative control (water with no stressor added). Optical density was measured every ten minutes for 24 hours after back-dilution using a Tecan Spark plate reader.

After 24 hours of growth, data from the plate reader was retrieved and the percent growth was calculated for each replicate using the average of the water controls in the same plate. Plates were collected and initially spun down at 3200 x *g* for 15 minutes before supernatants were placed into a 96-well filter plate to remove any remaining bacterial cells (Millipore Sigma #MAGVS2210) and spun at the same speed for 15 minutes until the supernatants have completely passed through the filter. Percent growths were calculated by normalizing endpoint ODs in the same plate to the average of the water controls.

Statistical significance was determined by ANOVA with a post-hoc Tukey’s HSD test, testing for whether any of the treatments significantly differed from water controls within the same plate.

### Lysate normalizations and qPCR to quantify prophage induction

Supernatants from treatments where the percent growth of the culture was less than 25% of the no-treatment controls were excluded from further screening. The resulting supernatants were normalized by final OD before adding DNase (Norgen Biotek #25710) to each well to remove any DNA that is not encapsulated by phage capsids. Supernatants were incubated for 15 minutes at room temperature before the DNase was denatured at 75 °C for 5 minutes.

One or two µL of DNase-treated supernatant were used as templates for qPCR. Primers for each prophage were designed to amplify a 200 – 300 bp region spanning the *attP* sites computed by the TIGER/Islander software (Mageeney *et al*. 2020b) (**Supplemental Table S5**). Amplification of the *attP* site indicates excised and circularized phage genomes, a proxy for prophage activity in our samples. Each well of the supernatants was tested in triplicate in 384-well plates (Fisher Scientific #14-380-055) as 10 µL qPCR reactions with the Luna® Universal qPCR Master Mix (New England Biolabs #M3003L) following the manufacturer’s directions. -ΔCt values were calculated for each treatment, where -ΔCt = –(ΔCt_treatment_average_ – ΔCt_water_average_) within the same plate. Treatments with statistically significant impacts were determined via ANOVA with a post-hoc Tukey’s HSD test, comparing Ct values of the different treatments to water controls. Significant treatments are indicated in **Supplemental Table S6**.

## Acknowledgements

We would like to thank Hazel Sisson and the members of the Phage Foundry (https://phagefoundry.org/) for their thoughtful feedback on the manuscript. J.T., V.K.M., and C.M.M. conceived this study. J.T. performed all the assays, with help from H.K.C. for the initial EC50 determinations and experimental design. All authors contributed to the writing and editing of the manuscript.

This material by the Biopreparedness Research Virtual Environment (BRaVE) Phage Foundry at Lawrence Berkeley National Laboratory is based upon work supported by the U.S. Department of Energy, Office of Science, Office of Biological & Environmental Research under contract number DE-AC02-05CH11231.

Sandia National Laboratories is a multi-mission laboratory managed and operated by National Technology & Engineering Solutions of Sandia, LLC (NTESS), a wholly owned subsidiary of Honeywell International Inc., for the U.S. Department of Energy’s National Nuclear Security Administration (DOE/NNSA) under contract DE-NA0003525. This written work is authored by an employee of NTESS. The employee, not NTESS, owns the right, title and interest in and to the written work and is responsible for its contents. Any subjective views or opinions that might be expressed in the written work do not necessarily represent the views of the U.S. Government. The publisher acknowledges that the U.S. Government retains a non-exclusive, paid-up, irrevocable, world-wide license to publish or reproduce the published form of this written work or allow others to do so, for U.S. Government purposes. The DOE will provide public access to results of federally sponsored research in accordance with the DOE Public Access Plan.

