## Supplemental Figure for "Chemical stressors differentially induce prophage replication in *Acinetobacter baumannii* strains"

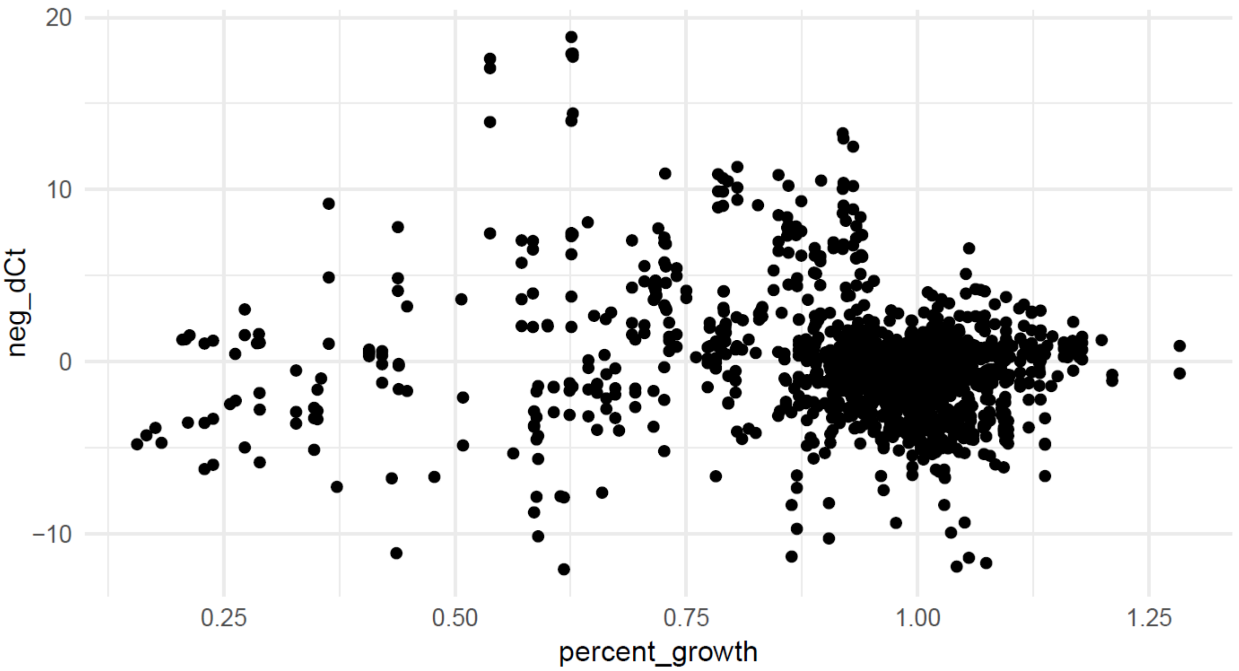


**Supplemental Figure S1. Scatter plot correlating the percent growth of *A. baumannii* in growth experiments to the –ΔCt values of their respective prophages from qPCR assays.** A –ΔCt value around 0 represents no difference in prophage replication relative to water controls, and a percent growth around 1.00 represents no difference in bacterial growth relative to water controls. Note that –ΔCt values were not evaluated for any *A. baumannii* conditions where percent growth was below 0.2.

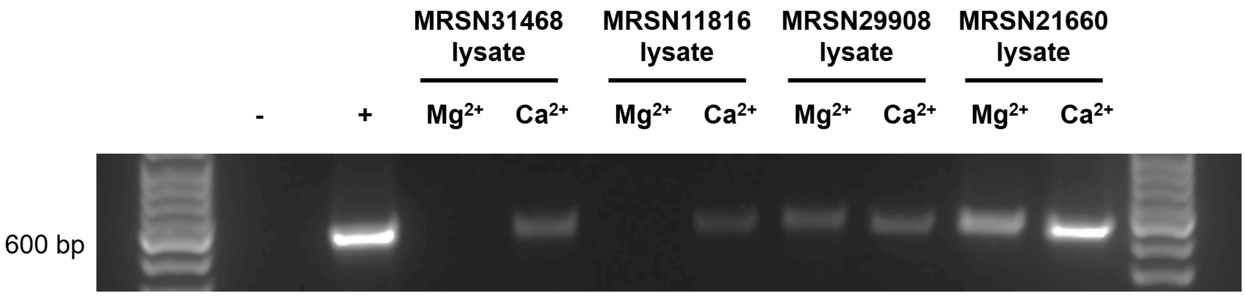


**Supplementary Figure S2. Agarose gel showing 16S PCR products amplified from lysates of tested strains.** Lysates before normalizations and DNase treatment were used as templates for each reaction so that the concentration of magnesium and calcium ions are the same across each sample. Note that the growth of MRSN31468 and MRSN11816 are significantly inhibited in magnesium and may contribute to the lack of a visible band for 16S.

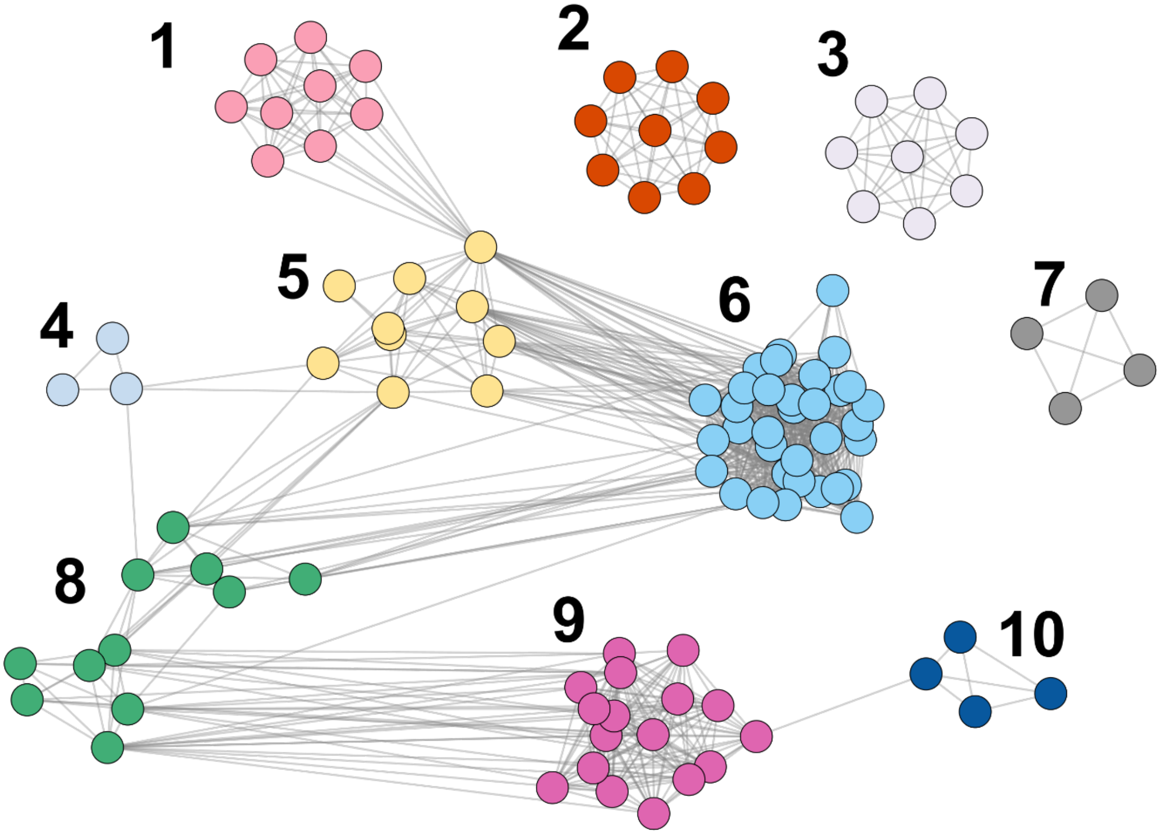


**Supplemental Figure S3. Cluster map of the prophages predicted from the *Acinetobacter baumannii* genomes.** Clusters are determined by amino acid identity and percent of genes shared.
